# Are signals of aggressive intent less honest in urban habitats?

**DOI:** 10.1101/504258

**Authors:** Çağlar Akçay, Michelle L. Beck, Kendra B. Sewall

## Abstract

The effect of urban noise on animal communication systems is one of the best examples of how anthropogenic change affects animal social behaviour. Urban noise often drives shifts in acoustic properties of signals but the consequences of noise for the honesty of signals – that is, how well they predict signaler behaviour, is unclear. Here we examine whether honesty of aggressive signaling changes in urban living song sparrows (*Melospiza melodia*). Aggressive signaling in song sparrows consists of close-range signals in two modalities that predict a subsequent attack: the low amplitude soft songs (an acoustic signal) and wing waves (a visual signal). Male song sparrows living in urban habitats display more territorial aggression than males living in rural habitats, but whether the honesty of close-range signals is affected by urbanization has not been examined. If soft songs are less effective in urban noise, we predict that they would be less reliably associated with attack in these habitats compared to rural habitats. We found that while acoustic noise was higher in urban habitats, the urban birds still sang more soft songs than rural birds during a simulated territorial intrusion. Furthermore, high rates of soft songs and low rates of loud songs predicted attacks in both habitats. Finally, we found evidence for a potential multimodal shift: urban birds tended to give proportionally more wing waves than soft songs than rural birds. These results indicate that urbanization might have a limited effect on the overall honesty of aggressive signals in song sparrows.

## Introduction

When individuals with conflicting interests communicate (e.g. during an aggressive interaction) there is an incentive for each signaler to try to manipulate the receiver into behaving in a way that benefits the signaler, thus jeopardizing the honesty of the signal (Dawkins and Krebs, 1978). This problem is particularly pronounced for signals of aggressive intent which are by definition not tied to a physical trait of the signaler. Instead they are thought to predict future behavior of the signaler. These signals are usually not costly to produce and can potentially be given at any level. A good example of this is bird song: singing seems to carry little or no metabolic cost compared to other activities birds have to carry out during an aggressive interaction (Zollinger et al., 2011).

Although early ethological literature assumed these signals of intent had to be honest – otherwise they would not exist – theoretical and empirical treatments of these signals in the 1970s were more skeptical (Caryl, 1979; Dawkins and Krebs, 1978; Maynard Smith, 1974). The problem seemed to be that if signals are only indicative aggressive “intent” of the signaler and not tied to a physical cost, then the signals would be easy to cheat for “bluffers” who would threaten without any intention to follow through with an attack. Therefore these signals were viewed mostly as attempts at manipulation by the signaler instead of carrying information regarding future behavior (Dawkins and Krebs, 1978). More recently, however, a multitude of studies have shown that signals of aggressive intent can honestly predict a subsequent escalation such as an attack in many species (e.g. Akçay et al., 2013; Bachmann et al., 2017; Laidre, 2009; Searcy et al., 2006; Waas, 1991). Often, the mechanism that ensures the honest of these signals seem to be the subsequent risk of retaliation from receivers (Anderson et al., 2013; Anderson et al., 2012; Bachmann et al., 2017; Molles and Vehrencamp, 2001).

An implicit assumption in the studies of honest signaling has been that the signaling systems are at an evolutionary equilibrium such that signaling strategies persist over non-signaling strategies (Searcy and Nowicki, 2005). Changes in physical and social ecology however, may disrupt this equilibrium. One such change that animal populations currently experience is human-induced environmental change, in particular urbanization (Johnson and Munshi-South, 2017). Although there are a plethora of studies on the effect of urbanization on signal features, particularly with respect to acoustic noise and song (Brumm and Slabbekoorn, 2005; Derryberry et al., 2016; Gil and Brumm, 2014; Halfwerk and Slabbekoorn, 2009; Patricelli and Blickley, 2006; Wood and Yezerinac, 2006) how the overall honesty of signaling systems change is less well studied, particularly in aggressive signaling.

Several studies showed that birds living in urban and rural habitats exhibit significant differences in responses to simulated territorial intrusions, with urban birds responding more strongly to simulated territory intrusions than rural birds (Davies and Sewall, 2016; Evans et al., 2010; Fokidis et al., 2011; Foltz et al., 2015; Hardman and Dalesman, 2018). However, these studies did not determine if aggressive signals also differed in their honesty between habitats. It is worth noting that although honest aggressive signals are correlated with other aggressive behaviours like approaching and attacking an opponent, these signals (unlike approach and attack) have no physical function in the aggressive interaction other than the information they carry (Otte, 1974). Thus, aggressive signals and non-signaling aggressive behaviours constitute separate behavioural characters and may respond differently to changes associated with urbanization (Akçay et al., 2015b; see Araya-Ajoy and Dingemanse, 2014 for a discussion of behavioral characters). To our knowledge no previous study assessed the honesty of aggressive signals in urban and rural habitats.

Here we ask whether the honesty of multi-modal signals of aggressive intent differs between urban and rural male song sparrows, *Melospiza melodia*, a songbird common in North America and found abundantly in urban and rural habitats. Urban song sparrows have been found to exhibit higher levels of aggression than their rural counterparts in several studies (Davies and Sewall, 2016; Evans et al., 2010; Foltz et al., 2015). Song sparrows have a well-studied aggressive signaling system that consists of two close-range aggressive signals: low amplitude “soft” songs and wing waves (rapid fluttering of one or both wings without getting airborne) both of which predict a subsequent attack (Akçay et al., 2013; Nice, 1943; Searcy et al., 2014; Searcy et al., 2006). Loud (broadcast) songs however, do not reliably predict attack in this species (Searcy et al., 2014). This difference in honesty between soft songs and loud songs seems to hold for several other species: where soft vocalizations reliably predict attack (Akçay et al., 2015a), but loud vocalizations do not (Searcy and Beecher, 2009).

Soft songs and wing waves present an interesting potential case of how multi-modal signaling changes in urban habitats (Halfwerk and Slabbekoorn, 2015). The defining feature of these signals is the low amplitude compared to the loud broadcast songs which may be an adaptation to reduce transmission distances. In some species, soft song also differs in acoustic structure from broadcast songs (Dabelsteen et al., 1998; Vargas-Castro et al., 2017) although whether these differences are adaptations to decrease transmission distances further is currently unclear (Akçay and Beecher, 2012; Vargas-Castro et al., 2017). The low amplitude of the signal along with potential acoustic adaptation to decrease transmission distances would make soft songs less effective compared to louder signals due to the masking effect of high anthropogenic noise levels commonly found in urban habitats (Pohl et al., 2009). In the case of song sparrows in particular, Wood and Yezerinac (2006) found that most of the acoustic noise in urban habitats was present at 1-4 kHz range and that urban song sparrows living in noisy habitats put relatively less energy into this frequency range of their songs when singing loud songs. Soft song differs from loud song in song sparrows in that it has a lower minimum frequency (1500 to 1700 Hz for soft songs vs. ca. 2000 for loud songs, Anderson et al., 2008). Furthermore, in rural habitats birds tend to put relatively more energy into the lower frequencies of soft song which overlap with urban noise (Anderson et al., 2008). Thus, soft song may be particularly prone to interference from urban noise.

One solution to the presence of urban noise is to sing loudly. Indeed, animals often respond to noise by vocalizing at higher amplitudes in response to higher noise levels, which is termed the Lombard effect (Brumm, 2004; Brumm and Todt, 2002; Brumm and Zollinger, 2011; Cynx et al., 1998). The Lombard effect is particularly strong if noise overlaps the frequency range of the vocalizations (Brumm and Todt, 2002; Manabe et al., 1998). If song sparrows show a Lombard effect in urban areas, they may sing loud songs instead of soft songs to signal their aggressive intent. Under this prediction we expect more loud songs in the urban habitats compared to rural habitats particularly by those birds who end up attacking their opponent.

Another solution for the problem introduced by noise would be to close the distance to their opponent (the intended recipient of low amplitude vocalizations) in order to ensure transmission of low amplitude signals in the urban habitats (Halfwerk et al., 2012). There is evidence that birds are sensitive to the relationship between amplitude and distance to the receiver (Brumm and Slater, 2006b). Getting closer to the receiver during an aggressive interaction may come at a cost however, as the proximity to the receiver potentially increases the risk of retaliation (Anderson et al., 2012; Templeton et al., 2012).

A further strategy to ensure transmission of soft songs would be to increase repetition rate or serial redundancy (Brumm and Slater, 2006a). Under this strategy we expect the rate of soft songs to increase in urban habitats, while rates of loud songs should not change given the latter do not reliably signal aggression. These strategies (increasing the amplitude of soft songs, decreasing distance and increasing serial redundancy) are not mutually-exclusive strategies, however they would affect the overall honesty of the signal, measured as a statistical association between the signal and subsequent attack in different ways. If urban song sparrows increase the amplitude of their aggressive songs we expect that they would sing more loud songs compared to rural song sparrows, and attackers would give significantly more loud songs, making loud songs the more honest signal. If song sparrows decrease the distance to the mount while singing soft songs we expect the distance while singing softly will be lower in urban than rural habitats while distance while singing loud songs would not differ between urban and rural habitats. The latter prediction assumes that the intended audience of the loud songs is not the immediate intruder but other neighbors, since loud songs do not reliably predict attack on the immediate intruder. Finally, if song sparrows increase the repetition rates for soft songs in urban habitat, we expect birds will sing more soft songs in urban habitats, and this difference will be particularly pronounced for attackers.

Given that song sparrows also have a visual signal of aggression, wing waves, that is positively correlated with soft songs, urban song sparrows may also shift their signaling effort to the visual modality (Halfwerk and Slabbekoorn, 2015). Only a few studies have examined whether acoustic noise drives such a multi-modal shift to visual signals and evidence for this remains absent in birds (Grafe et al., 2012; Partan, 2017; Patricelli and Blickley, 2006; Ríos-Chelén et al., 2015). If urban song sparrows indeed switch to the visual modality, we might expect that they would give more wing waves and fewer soft songs. Previous studies in song sparrows reported strong positive correlations between the wing waves and soft songs (Akçay et al., 2014; Nice, 1943; Searcy et al., 2006). We therefore predicted that if urban birds increase their use of wing waves while decreasing their use of soft songs, the correlation between wing waves and soft songs should be absent or weaker in the urban birds compared to rural birds.

## Methods

### Study site and Subjects

We studied song sparrows in Montgomery County, VA at 2 urban and 3 rural sites. The two urban sites were the campuses of Virginia Tech (Blacksburg, VA) and Radford University (Radford, VA). The three rural sites were Heritage Park (just outside Blacksburg, VA), Kentland Farms of Virginia Tech and Stroubles Creek Stream Restoration area. These sites differ significantly in their urbanization based on quantitative measures of vegetation, paved surfaces, and buildings (Davies et al., 2018). Trials were carried out between 8^th^ April and 13^th^ May 2017. Most subjects tested were unbanded at the time of the trial but were captured after the simulated territory intrusion for banding and blood sampling for a different study. We tested 42 rural birds and 36 urban birds.

### Noise Measurements

We measured the ambient noise levels at a randomly selected subset of the territories in experiment (12 urban and 16 rural territories) during morning hours (0600 to 1200 hrs) using a sound meter (Radioshack Digitial Sound Level Meter model 33-2055) in setting A and fast response (125 ms) following the methods described in Brumm (2004). The A setting has a flat response within 1 to 8 kHz which covers most of song sparrow song range. To take the measurements we pointed a sound meter, oriented horizontally, in one of the cardinal directions, picked randomly. We noted the maximum sound level measurement in a 10 second period and then rotated the sound meter clock-wise by 90 degrees and repeated the measurement. We took 2 measurements per cardinal direction and then averaged the eight values. Although this method does not quantify noise in specific frequency ranges, it has been shown that noise measured in this way is functionally relevant to singing behaviors in several species (e.g. Brumm and Slater, 2006a).

### Song stimuli

We recorded songs from male song sparrows around Blacksburg and Radford for making stimuli using a Marantz PMD 660 or PMD 661 Solid State recorder and a Sennheiser ME66/K6 directional microphone. From these recordings we chose the song types that had a high signal to noise ratio from the recordings. We used 38 song types from 24 different males during the experiment. The majority of the stimuli (24 out of 38) came from males holding territories in residential areas and parks in Blacksburg as well as the edge of campus where the habitat grades into fields. Two songs came from rural sites, and the rest came from Radford University and Virginia Tech Campuses. The stimuli for each subject came from a male that was at least 1 km away (in most cases more than 5 km) from that subject, thus representing an unfamiliar song. We never used a song recorded from the same site as a stimulus during a behavioral trial.

### Aggression assays

We carried out the simulated territory intrusions at a location that was estimated to be a central location in the male’s territory based on observation of singing perches. We placed a speaker (VictSing model C6 connected to a smartphone via Bluetooth) and a taxidermic model of a song sparrow on a natural perch that was initially covered by a cloth. We adjusted speaker volume to be approximately 80 dB SPL, measured at 1m (with the same sound meter and settings as above), which corresponds to loud song volume in song sparrows. Two observers standing about 20m from the speaker narrated the trial with the same recording equipment.

After setting up the equipment, we started to play a song at a rate of one song every ten seconds with the taxidermic model covered. Song sparrow songs last an average of 3 seconds and we presented stimuli at a rate of one song per 10 seconds for the duration of the trial which approximates typical song sparrow singing rate. Each male received only a single rendition of one song type during the trial repeated every 10 seconds. This is consistent with the fact that song sparrows repeat a single song type for several minutes during their natural singing (eventual variety singing), and does not lead to any habituation at even longer durations than used in this experiment (Akçay et al., 2013). We recorded behaviours for three minutes after the first response of the focal male (the pre-mount period). After the pre-mount period, we paused the playback and one experimenter removed the cover to reveal the taxidermic model. We then restarted the playback at the same song rate as before and continued for another 10 minutes or until the subject attacked, physically touched the mount, at which point we stopped the playback and retrieved the mount before it was destroyed (the mount period).

### Response Measures

During the trial, the observer narrating the trial noted attacks and the following behaviours: flights (with distance to the speaker after each flight), soft songs, loud songs and wing waves (all divided by trial duration and reported as rates). Soft and loud song determination was made in the field by experienced observers (CA or MLB). Song amplitude in song sparrows varies continuously between 55 dB to 85 dB, and our determination of soft vs. loud song reduces this continuous variation into a categorical decision. This method has been validated by Anderson et al. (2008) who showed that an expert observer produces a clear cut-off point with soft vs. loud determinations made in the field when these are validated with actual amplitude measurements from a fixed distance. Several studies using soft song categorization in this way in this species found that it reliably predicts attack whereas loud songs do not (Akçay et al., 2015a). Thus, this categorization captures biologically meaningful variation in amplitude.

The trial recordings were scanned with the software Syrinx (John Burt, Portland, OR). From the trial recordings we extracted the counts of flights, loud songs, soft songs and wing waves and proportion of time spent within 1m for both the initial pre-mount period and the mount period. Additionally, we noted the closest approach distance for the pre-mount period. We did not use closest approach for the mount period as a response variable because there was little variation in that measure for the mount period (an overwhelming majority of the subjects approached to within 1m). Flights, proportion of time spent within 1 m of the speaker, closest approach (pre-mount period) are considered aggressive behaviours, whereas the loud songs, soft songs and wing waves are considered signaling behaviours (Akçay et al., 2015b). Finally, we also extracted from the recordings the distance at which each loud and soft song were delivered (as noted above, distance information was given with each flight during the trial).

### Data analyses

Our first analysis addressed whether there were any differences between aggressive behaviours and signaling behaviours of rural and urban birds. We used Mann-Whitney U tests for all aggressive behaviors and signaling behaviours as these were non-normally distributed. We report effect sizes (Hedges’ g, computed with the R package “effsize”; Torchiano, 2018) and confidence intervals for the urban-rural comparisons in all of the response variables. We carried out a Chi-square test to determine whether attack rates differed between urban and rural birds.

To address our main question of whether honesty of signaling differs between urban and rural habitats, we carried out separate logistic regressions with attack as the dependent variable (attack or non-attack) and the following as the predictor variables: habitat and signal (soft songs, wing waves or loud songs), and the interaction between habitat and the signal. The main effects and interaction effects were entered sequentially, representing 2 contrasts we were interested in: 1) Does a signal (soft song, loud song or wing waves) predict attack after taking into account the effect of habitat and 2) Is there an interaction between habitat and signal in predicting attack? We also compared the proportion of soft songs among all songs of attackers and non-attackers in a similar logistic regression model. In supplementary materials (Tables S3-S6) we also report parallel analyses with general linear mixed models in which used the same fixed effects but also added site as a random factor. These results closely parallel the models reported in the main text but the models showed singular fits. We therefore report the models without the site as a random factor below.

To determine whether urban soft songs were given at a shorter distance from the speaker/mount we determined for each subject the average distance at which soft songs and loud songs were given, separately for both the pre-mount and mount periods. We then carried out a linear mixed model with habitat, type of song (soft vs. loud) and their interaction as fixed variables and subject as the random variable.

Finally, we asked whether there was a multi-modal shift from soft songs to wing waves in urban areas. Soft songs and wing waves are highly correlated with each other (Akçay et al., 2014). Thus, we need to control for the level of overall signaling effort to determine whether wing waves were more common in urban areas. In order to do that, we first added together all aggressive signals (counts of wing waves and soft songs) and then took the proportion of wing waves among the total number of aggressive signals for subjects who gave at least one soft song or wing wave. We then compared the proportion of wing waves between the habitats with a Mann-Whitney U test.

## Results

Ambient noise levels were significantly higher by approximately 8 decibels in the urban territories (M±SD: 71.22±3.11 dB; n=12) than the rural territories (M±SD: 64.37±5.54 dB, n=16; independent samples t-test: t_26_=3.84, p=0.0007). The noise levels at urban habitats correspond to the higher end of noise measurements reported in a study that documented effects of urban noise on the acoustic properties of song sparrow song (Wood and Yezerinac, 2006). Urban birds were significantly more aggressive than rural birds in all of the aggressive behaviors except rate of flights during the mount period (Table 1 and Table 2). More urban birds (14 out of 36, 38.9%) attacked the mount than rural birds (4 out of 42, 9.5%; χ^2^= 9.42; p= 0.002).

**Table 1.** Comparison of urban and rural song sparrows in the response variables during the pre-mount period. Positive effect sizes mean higher values for urban birds.

| Variable | W | p-value | Hedges' g | 95% CI |
| --- | --- | --- | --- | --- |
| *Flight rate* | 500 | 0.0099 | 0.59 | 0.13 – 1.06 |
| *Closest approach* | 1129 | 0.000002 | -0.56 | -0.10 – 1.02 |
| *Proportion of time within 1m* | 376.5 | 0.00013 | 0.95 | 0.47 – 1.42 |
| *Soft song rate* | 397 | 0.00028 | 0.69 | 0.23 – 1.16 |
| *Loud song rate* | 645 | 0.27 | 0.19 | -0.26 – 0.65 |
| *Wing wave rate* | 385 | 0.000008 | 0.78 | 0.31 – 1.25 |

**Table 2.** Comparison of urban and rural song sparrows in the response variables during the mount period. Positive effect sizes mean higher values for urban birds.

| Variable | W | p-value | Hedges' g | 95% CI |
| --- | --- | --- | --- | --- |
| *Flight rate* | 624 | 0.19 | 0.39 | 0.06 – 0.86 |
| *Proportion of time within 1m* | 358.5 | 0.000007 | 0.93 | 0.45 – 1.40 |
| *Soft song rate* | 558 | 0.046 | 0.50 | 0.04 – 0.96 |
| *Loud song rate* | 825.5 | 0.49 | -0.06 | -0.51 – 0.39 |
| *Wing wave rate* | 524 | 0.015 | 0.38 | -0.07 – 0.83 |

During the pre-mount period, urban birds sang more soft songs and gave more wing waves than rural birds. Loud song rates did not differ significantly between urban and rural birds during the pre-mount period (Table 1). During the mount period, urban birds sang more soft songs and gave more wing waves than rural birds. Loud song rates did not differ significantly between urban and rural birds during the mount period (Table 2).

Logistic regression models on attacks as the response variable showed that the main effect of habitat and each of soft song (Figure 2a), wing wave (Figure 2b) and loud song (Figure 2c) was significant (Table 3): Birds that sang high rates of soft songs, gave high rates of wing waves and were more likely to attack. Interestingly, birds that sang fewer loud songs were also more likely to attack. Consequently, the proportion of soft songs was also a highly significant predictor of attack: birds that sang a higher proportion of soft songs were more likely to attack (Figure 2d, Table 3). The two-way interaction between the signal and habitat was not significant in any of the models, suggesting the honesty of signaling did not differ between urban and rural habitats

**Table 3.** Logistic regression models with habitat and soft songs, wing waves, loud songs or proportion of soft songs during the mount period as predictor variables. A separate model for each signal was run. The cells report χ^2^ values (p-values, alpha <0.05 indicated with bold text) from a forward sequential logistic regression. Note that in all models, we entered habitat first, followed by the signal and the interaction term. We excluded six subjects that did not sing any songs (soft or loud) from the model with proportion of soft songs (rightmost column).

| Model: | *Soft song model* | *Wing wave model* | *Loud song model* | *Proportion of soft songs* |
| --- | --- | --- | --- | --- |
| **Habitat** | 9.74 (**0.002**) | 9.74 (**0.002**) | 9.74 (**0.002**) | 5.18 (**0.022**) |
| **Signal** | 5.13 (**0.023**) | 5.36 (**0.020**) | 17.28 (**0.000003**) | 13.21 (**0.0003**) |
| **Habitat\*Signal** | 0.32 (0.57) | 2.43 (0.12) | 1.14 (0.28) | 0.65 (0.42) |

**Figure 1.**
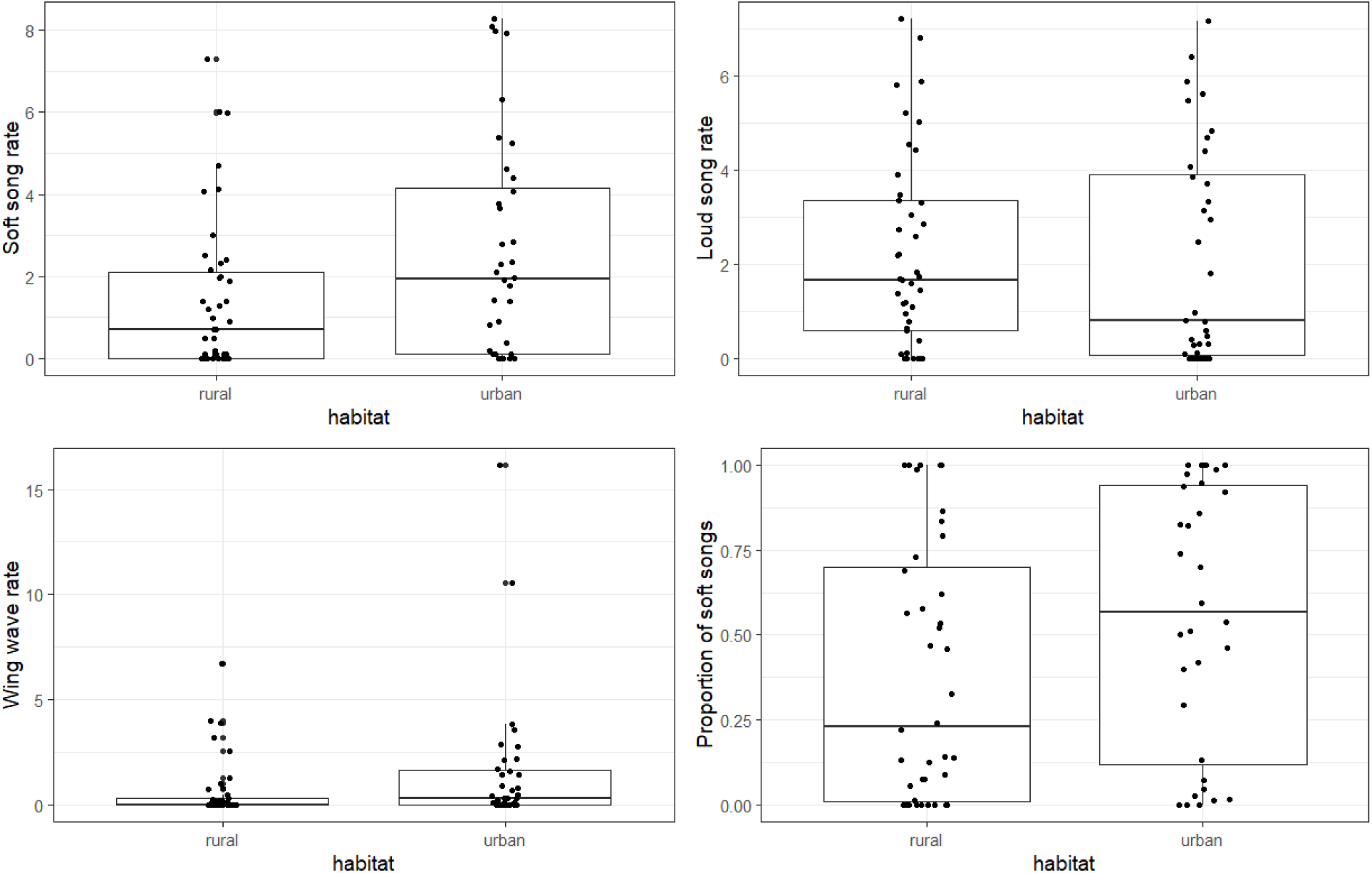
Mount period (a) soft song rates (b) loud song rates, (c) wing wave rates and (d) proportion of soft songs. The dots are individual data points, boxes indicate the interquartile range and medians. Whiskers are 95% confidence intervals. Rates are given per minutes.

**Figure 2.**
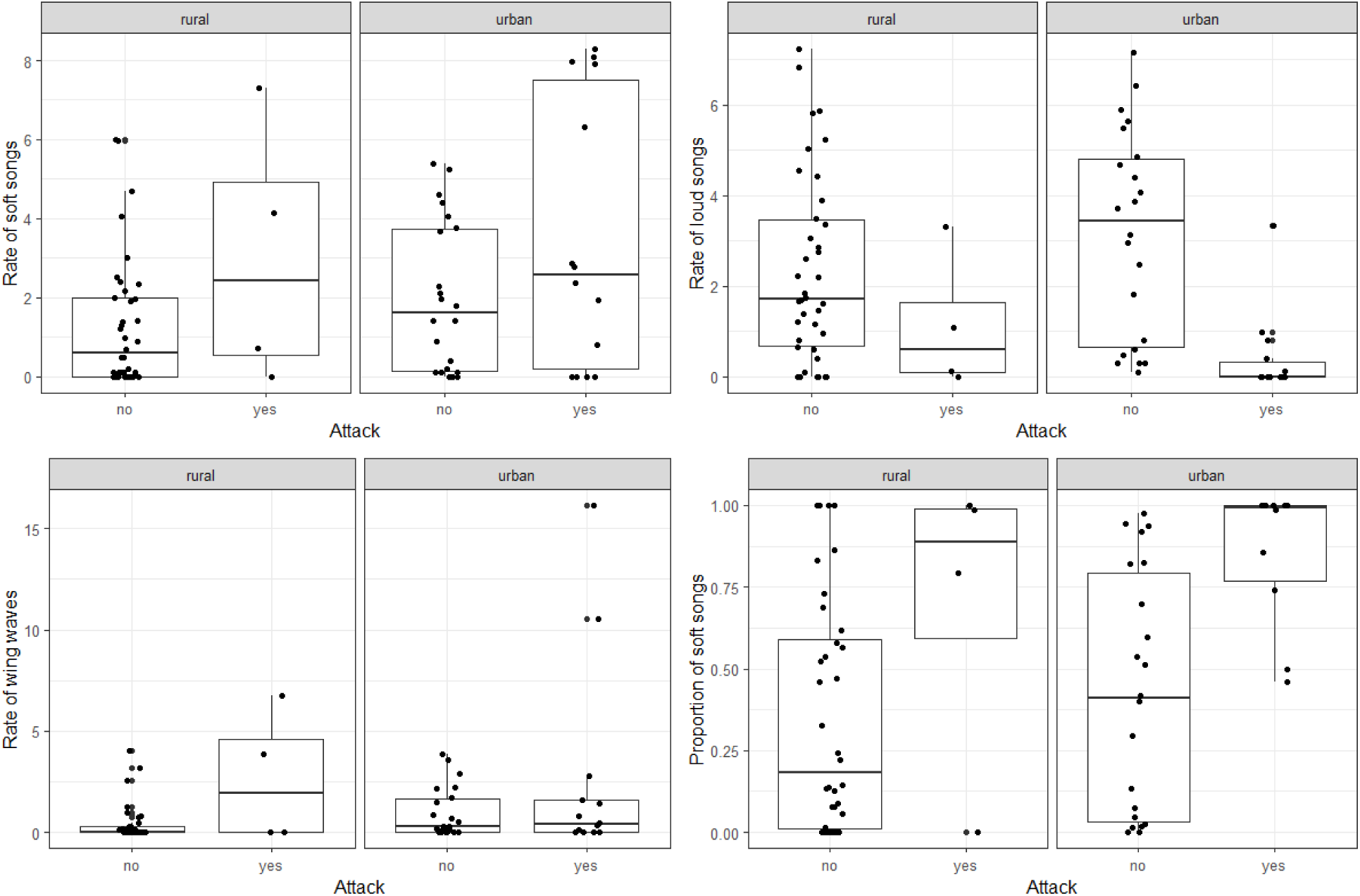
Mount period (a) soft song rates, (b) wing wave rates, (c) loud song rates and (d) proportion of soft songs of attacking and non-attacking birds by habitat. The dots are individual data points, boxes indicate the interquartile range and medians. Whiskers are 95% confidence intervals. Rates are given per minutes.

In general, soft songs were sung closer to the speaker than loud songs for both urban and rural birds. Furthermore, urban birds sang both soft and loud songs closer to the speaker than rural birds (Table 4, Figure 3). The linear mixed model on song distances for loud and soft songs showed a significant effect of habitat and song category (soft songs were sung in closer proximity to the speaker) but no interaction effect between habitat and song category (Table 5).

**Table 4.** Means (SD) of song distances in rural and urban habitats for loud songs and soft songs in the pre-mount and mount periods.

|  | Pre-mount |  | Mount |  |
| --- | --- | --- | --- | --- |
|  | rural | urban | rural | urban |
| **loud songs** | 6.14 (4.16) | 3.99 (2.83) | 5.61 (3.92) | 3.71 (3.95) |
| **soft songs** | 4.40 (3.71) | 2.15 (3.06) | 1.92 (2.67) | 1.06 (2.13) |

**Table 5.** Linear mixed models on average distances depending on habitat (urban vs rural) and song category (soft vs loud) during the pre-mount and mount periods.

| *Pre-Mount period* | *Coefficient (SE)* | *t* | *p* |
| --- | --- | --- | --- |
| **Intercept** | 6.01 (0.59) | 10.24 | **<0.00001** |
| **Habitat (urban)** | -2.14 (0.85) | -2.51 | **0.014** |
| **Song category (soft)** | -1.41 (0.65) | -2.17 | **0.035** |
| **Habitat\* Song category** | -0.20 (0.90) | -0.23 | 0.82 |
| *Mount Period* |  |  |  |
| **Intercept** | 5.48 (0.54) | 10.05 | **<0.00001** |
| **Habitat (urban)** | -1.88 (0.83) | -2.27 | **0.026** |
| **Song category (soft)** | -3.13 (0.56) | -5.59 | **<0.000001** |
| **Habitat\* Song category** | 0.78 (0.82) | 0.96 | 0.34 |

**Figure 3.**
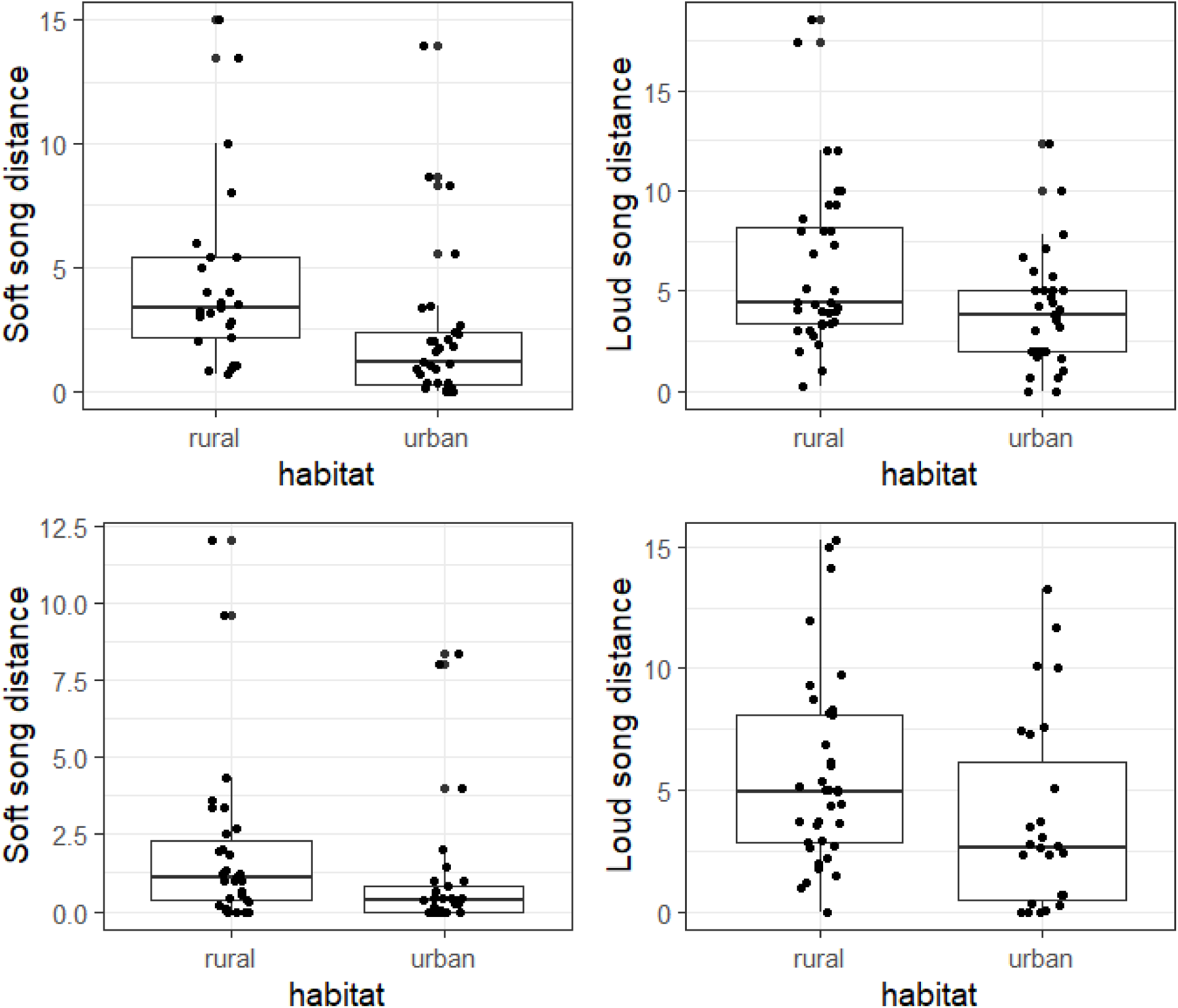
Song distances during the pre-mount (a, b) and mount (c,d) periods. The dots are individual data points, boxes indicate the interquartile range and medians. Whiskers are 95% confidence intervals. The left panels show distances at which soft songs are sung, the right panels show distances at which loud songs are sung.

During the pre-mount period, there was a non-significant trend for urban birds to give proportionally more wing waves than rural birds (0.43 vs. 0.30 for urban vs. rural subjects) (U=348.5, p=0.074, n= 62, Hedges’ g= 0.39; 95% CI: −0.12–0.91). During the mount period, urban birds also gave proportionally more wing waves than rural birds (0.32 vs. 0.19), and the difference was significant (U= 302.5, p=0.043, n= 59, Hedges’ g= 0.49; 95% CI: −0.03–1.02; Figure 4).

**Figure 4.**
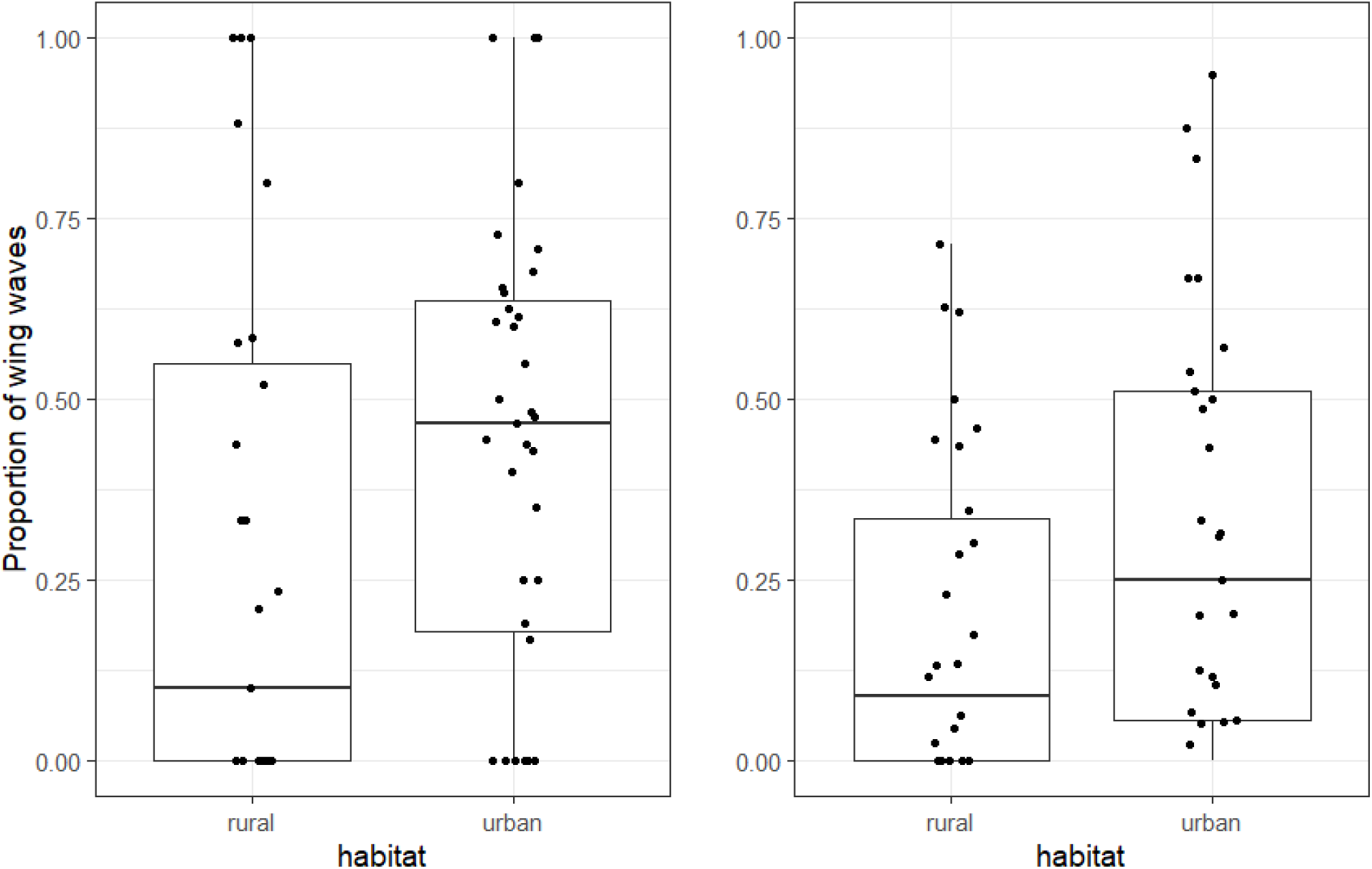
Proportion of wing waves among the sum of soft songs and wing waves for (a) pre-mount period and (b) mount period. The dots are individual data points, boxes indicate the interquartile range and medians. Whiskers are 95% confidence intervals.

## Discussion

We aimed to test the hypothesis that low amplitude songs in urban song sparrows may be a less honest signal of aggression than loud songs due to anthropogenic noise in urban habitats. Contrary to this prediction, soft songs were predictive of a physical attack in urban and rural habitats alike and song sparrows in urban habitats sang more soft songs than rural birds, consistent with the fact that they were also more aggressive than rural birds. Wing waves also showed the same pattern: urban birds gave more wing waves, again consistent with the fact they were more aggressive than rural birds. Wing waves also reliably predicted attack in both habitats. Interestingly, the most honest signal of attack in both habitats was low rates of loud songs: attackers sang fewer loud songs per minute than non-attackers. We found that urban birds generally sang at a shorter distance from the speaker for both loud and soft songs, and soft songs were given at a closer distance than loud songs in both habitat types. Finally, we found that during the mount period urban birds gave proportionally more wing waves as part of their total signaling effort. To our knowledge, this is the first study to examine honest multi-modal signaling in aggressive interactions in relation to urbanization and adds to the growing literature on behavioral effects of urbanization on animal social behavior. Below we discuss these results in the context of previous research on signal honesty in this and other songbirds.

### Song sparrows sing softly and closely in the city

Even under the noisy conditions of the urban habitats, male song sparrows seemed to use soft songs as an honest signal of aggressive intent. Given the low amplitude of soft song (relative to loud song), it would be reasonable to suppose that soft song will be less effective in urban habitats as an aggressive signal. The present results do not support this hypothesis. Instead, urban birds sang more soft songs than rural males. Increasing the rate of repetition and therefore the serial redundancy (Brumm and Slater, 2006a) may be a strategy to ensure the reception of the signal under noise, although it is worth noting that the rate of wing waves also increased in urban habitats compared to rural habitats. Given that wing waves, a visual signal, are not masked by acoustic noise this finding suggests that the increase in soft song rates in urban habitats may be due to the increased aggression levels of song sparrows.

A potential strategy that song sparrows might engage in to deal with urban noise is to close the distance to the receiver (Halfwerk et al., 2012) such that the signal-to-noise ratio of the acoustic signals would be improved at the receiver end of the transmission. Indeed, we found that urban birds sang at shorter distances to the speaker (the presumed receiver of the signals) for both loud songs and soft songs compared to rural birds. Approaching closer to the speaker would also mean that the playback songs, which were played at the same amplitude in urban and rural habitats, would also not suffer from decreased signal to noise ratios in urban habitats compared to rural habitats at the point of the reception.

The clearest example of such a spatial strategy in comes from an elegant experiment by Halfwerk and colleagues (2012). In this experiment, male great tits (*Parus major*) singing to their mates adjusted their singing locations to be closer to the nest box when they experimentally presented noise inside the nest box when their mates was in the nest box. Remarkably, the males did not experience the noise themselves (as the noise was only presented in the nest box and was not audible outside) but evidently acquired the information about the noise socially from their mates. In another recent study, male white-crowned sparrows living in noisier territories approached the speaker closer than the males in the same population that lived in quieter territories. (Phillips and Derryberry, 2018). One interpretation of this finding is that high levels of ambient noise might require birds to approach each other closer to evaluate and transmit signals efficiently. A similar logic may apply to song sparrows in our urban habitats as well. Unlike the Halfwerk et al. study, however, neither the current study nor Phillips and Derryberry (2018) experimentally manipulated noise levels to allow a causal inference about the role of noise in determining proximity in aggressive interactions. Interestingly, in another experimental study, European robins (*Erithacus rubecula*) were found to move away from a source of noise as the volume of the noise increased, although the noise presentation in that case was not simultaneously accompanied with song stimulus (McLaughlin and Kunc, 2013). Therefore, the males did not have a reason to stay in close proximity to the source of noise.

### Multimodal signaling in urban habitats

Although urban noise did not decrease the use or honesty of song songs we found tentative evidence for a multi-modal shift: urban birds tended to give more wing waves proportionally to their total aggressive signaling effort at least during the mount period although the effect size was moderate and the confidence intervals were large. This finding, if confirmed, is consistent with the hypothesis that acoustic noise found in urban habitats may lead to switching signaling effort to the visual modality (Partan, 2017; Partan et al., 2010). It is also important to note that if a multi-modal shift is occurring in urban song sparrows it is incomplete: the urban birds still sing more soft songs than rural birds and soft song is still an honest signal of aggressive intent in urban birds.

Whether wing waves are a more effective signal compared to soft songs in urban habitats (compared to rural habitats) is an open question. To determine the relative effectiveness of these signals an experiment displaying the visual (wing wave) and acoustic (soft song) signal separately and together with a robotic model would be required (Anderson et al., 2013; Partan et al., 2010; Partan et al., 2009). To the best of our knowledge only one experiment compared responses to signals in different modalities in urban and rural habitats. In this study, Partan and her colleagues (2010) found that urban gray squirrels (*Sciurus carolinensis*) responded more to the visual alarm signal (tail flagging) displayed by a robotic squirrel than the rural squirrels. There was however no significant difference in response strength to the vocal signals between urban and rural squirrels. These results suggest that urban gray squirrels may rely more on the visual signals in urban habitats even if vocal signals are still as effective in urban habitats as in rural habitats.

In another relevant study Ríos-Chelén and colleagues examined whether red-winged blackbirds changed their signaling effort from acoustic to visual signals (the “song-spread display”, in which singing males spread their wings to expose their red epaulets) in noisier habitats (Ríos-Chelén et al., 2015). They found no effect of the ambient noise on the intensity of visual displays although males in the noisier habitats did change some features of their vocalizations.

How animals deal with noise in multiple modalities has been examined in relatively few studies, although it is increasingly becoming a focus of attention (Brumm and Slabbekoorn, 2005; Halfwerk and Slabbekoorn, 2015). We believe the aggressive signaling system of song sparrows (and related species like swamp sparrows; Anderson et al., 2013; Ballentine et al., 2008) provides an excellent model system to address how noise affects multi-modal signaling. As noted, wing waves and soft songs are highly correlated with each other and are therefore likely to be redundant, although noisy conditions in one modality may change the perception of these signals (Halfwerk and Slabbekoorn, 2015). Furthermore, the low amplitude of soft songs make it particularly likely to be prone to interference which may call for not only multi-modal shifts but also an increase in redundancy in signaling (e.g. a tighter correlation between wing waves and soft songs). These possibilities can be examined with experimental manipulations of ambient noise and multimodal signaling.

In summary, we found that urban song sparrows use soft songs as an honest signal, despite the expectation that urban noise may make it a less effective signal. Given the scarcity of studies on the honesty of acoustic signaling in urban habitats (despite a plethora of studies on how anthropogenic noise affects signal feature), it is still an open question whether urbanization in general alters honesty of communication systems found in less disturbed habitats. We also found that urbanization may affect multi-modal displays by inducing some males to switch to a visual display (wing waves) instead of soft songs. We believe that the song sparrow signaling system is an excellent model to ask how multi-modal signaling evolves under anthropogenic habitat change.

