## Supplemental Tables for "Are signals of aggressive intent less honest in urban habitats?"

Supplementary Tables and Figure- Akcay et al, submitted.

Table S1. Pearson correlation coefficients (p-values) for the variables in principle component analysis for pre-mount period n=78

|  | Proportion of time within 5m | Closest Approach |
| --- | --- | --- |
| Flights | 0.45 (p= 0.0000037) | -0.44 (p= 0.0000069) |
| Proportion of time within 5m | - | -0.65 (p< 0.00000001) |

Table S2. Pearson correlation coefficients (p-values) for the variables in principle component analysis for mount period n=78

|  | Proportion of time within 1m | Latency to 5m |
| --- | --- | --- |
| Flights | 0.56 (p< 0.0000001) | -0.38 (p= 0.00056) |
| Proportion of time within 1m | - | -0.47 (p= 0.000002) |

Table S3-6: results of general linear mixed models with the signaling variables and habitat as fixed factors and site as random factor. The response variable is attack (yes or no). The χ2 values are from a Wald-test. Note that the models had singular fits.

Table S3.

| **Soft song rates** | Chisq | Df | Pr(>Chisq) |
| --- | --- | --- | --- |
| **habitat** | 5.6461 | 1 | 0.01749 |
| **MountSoftsong** | 5.0605 | 1 | 0.02448 |
| **habitat:MountSoftsong** | 0.3173 | 1 | 0.57326 |

Table S4.

| **Wing wave rates** | Chisq | Df | Pr(>Chisq) |
| --- | --- | --- | --- |
| **habitat** | 5.125 | 1 | 0.02358 |
| **MountWingwave** | 4.123 | 1 | 0.0423 |
| **habitat:MountWingwave** | 2.3004 | 1 | 0.12934 |

Table S5.

| **Loud song rates** | Chisq | Df | Pr(>Chisq) |
| --- | --- | --- | --- |
| **habitat** | 7.2651 | 1 | 0.007031 |
| **MountLoudsong** | 6.6146 | 1 | 0.010114 |
| **habitat:MountLoudsong** | 1.1923 | 1 | 0.274866 |

Table S6.

| **proportion of soft songs** | Chisq | Df | Pr(>Chisq) |
| --- | --- | --- | --- |
| **habitat** | 2.3818 | 1 | 0.122757 |
| **ProportionSoft** | 8.1858 | 1 | 0.004222 |
| **habitat:ProportionSoft** | 0.6417 | 1 | 0.423098 |
